# Reproducible *in vivo* applications of the p16-3MR model in senescence research

**DOI:** 10.64898/2026.01.28.701695

**Authors:** Boshi Wang, Pilar Picallos Rabina, Jamil Nehme, Marco Demaria

## Abstract

The p16-3MR mouse model has been widely used to visualize and conditionally eliminate p16-expressing senescent cells *in vivo* and has been applied across diverse biological contexts, including tissue repair, fibrosis, cancer, therapy response, and aging. Despite the development of multiple senescence reporter and ablation systems, p16-3MR stands out for its broad and sustained adoption across independent laboratories. Here, we review the extensive published literature supporting the reproducibility and utility of the p16-3MR system and present new validation data across acute and chronic senescence-inducing conditions. We demonstrate reproducible induction of p16-associated bioluminescence during wound healing, chemotherapy, and aging, as well as partial but consistent reduction following ganciclovir treatment. We further delineate the strengths and limitations of the individual components of the 3MR construct, including HSV-thymidine kinase–mediated clearance, Renilla luciferase– based bioluminescence, and monomeric red fluorescent protein, and discuss how factors such as cell abundance, tissue context, pigmentation, and substrate chemistry influence detection sensitivity.

Together, these data confirm that the p16-3MR model is a functional and versatile tool for studying senescent cells *in vivo* when used with appropriate experimental design and interpretation, and they provide support for its continued application alongside emerging senescence models.

## Introduction

The p16-3MR mouse model has been broadly employed to study cellular senescence *in vivo*, enabling both the visualization and clearance of p16-expressing cells. Since its inception, several other reporter and clearance models of senescence have been proposed^1–8^. Yet very few have achieved the widespread adoption and breadth of applications seen with p16-3MR mice; the only other system with comparable use is the INK-ATTAC system^3^, which, like p16-3MR, couples a p16-driven suicide gene with reporter features to allow selective elimination of senescent cells. The extensive use of these systems across multiple independent groups reflects their continued relevance and reliability as cornerstone tools in senescence research. Importantly, studies have employed both the p16-3MR and INK-ATTAC models in parallel, and in these cases key findings have been reproduced across systems. For example, both models demonstrated delayed wound healing when p16-positive cells were ablated^3^.

Independent studies using the p16-3MR mice have reported induction of reporter signals associated with increased burden of p16-positive cells, efficient clearance of p16-positive cells upon ganciclovir administration, and corresponding functional effects across diverse biological contexts, including tissue repair, cancer, fibrosis, therapy response, and organismal aging^9–17^. Collectively, these studies demonstrate that the p16-3MR system remains a robust and versatile tool for dissecting the role of senescent cells in physiology and disease. Given that the original laboratory in which the model was developed is no longer active, our group has taken responsibility to assist colleagues who wish to use the p16-3MR mice, ensuring that the research community can continue to access and apply this model effectively.

As with all genetic reporter and ablation models, context-specific factors can influence signal strength and clearance efficiency. It is not unusual for transgenic models to display variability depending on breeding history, animal facility conditions, operators, or experimental equipment. Nevertheless, reproducibility has been established across multiple laboratories worldwide. Among the three components of the p16-3MR construct, the suicide gene has been the most widely and consistently used feature, followed by the luminescence reporter and, less frequently, the fluorescent signal. This pattern of use is not unusual and mirrors the experience with INK-ATTAC mice, where the suicide gene has been exploited extensively while the fluorescent component has been more limited in application.

The herpes simplex virus thymidine kinase (HSV-tk) component, which underlies the suicide function, is central to the model. Advantages include its strong track record in transgenic applications^18–20^, the high selectivity of ganciclovir-induced ablation, and the ability to eliminate both dividing and non-dividing p16^+^ cells, which is critical since most senescent cells are post-mitotic^21^. Moreover, clearance via HSV-tk/ganciclovir has been validated in multiple biological contexts, where elimination of senescent cells produced measurable functional effects. Disadvantages include possible incomplete ablation in tissues with low drug penetration or heterogeneous transgene expression, as well as potential off-target effects since ganciclovir can affect mitochondrial DNA in proliferating cells. Clearance efficiency also depends on the abundance of p16^+^ cells and on the expression levels of p16: smaller or low p16-expression populations may be more difficult to fully eliminate. Indeed, clearance efficiency is typically in the range of 50–75%, depending on tissue and experimental context, suggesting that not all p16^+^ cells can be effectively ablated, although the reasons for this remain to be fully understood.

The Renilla luciferase reporter^22,23^ offers complementary strengths and weaknesses. Advantages are its relatively small coding sequence, facilitating integration into complex constructs, its rapid light emission kinetics, and the possibility of dual-reporter applications alongside Firefly luciferase. Disadvantages are its lower brightness compared to Firefly luciferase^24^, requiring higher-abundance cell populations for robust detection, and greater sensitivity to substrate biodistribution, which can limit tissue penetration and *in vivo* sensitivity. In addition, coelenterazine substrate for Renilla is prone to auto-oxidation^25^, which can increase background signal; however, this is a well-known property of the chemistry and not specific to the p16-3MR system and careful controls and proper handling minimize this issue. Nevertheless, these features explain why luminescence from the p16-3MR system is reliable in contexts with abundant senescent cells, but less suited for detecting sparse populations.

Similarly, the monomeric red fluorescent protein (mRFP) component has both benefits and limitations. Advantages are that mRFP is small, monomeric (reducing aggregation), and relatively stable against photobleaching, making it useful for visualizing senescent populations in tissue sections or cultures. Disadvantages include relatively low quantum yield and brightness compared to newer fluorescent proteins, and a strong dependence on cell abundance and tissue context for detection. Consequently, while valuable under specific conditions, mRFP has been less widely used than the suicide or luciferase features.

The field of senescence research is still rapidly evolving, and we fully anticipate that improved models will be developed in the future. These may incorporate brighter reporters, enhanced clearance mechanisms, or humanized regulatory elements that better capture the complexity of senescence *in vivo*. We view the development and comparison of multiple models as a strength of the field, and the p16-3MR mice have played a critical role in establishing the foundation for these ongoing innovations.

## Results

In addition to the extensive independent literature summarized above, to further evaluate the reproducibility and functionality of the p16-3MR system, we conducted a series of validation experiments across different senescence-inducing contexts, ages, and sexes.

First, we confirmed reporter activation in well-established acute models. Following skin wounding with a 6-mm biopsy punch, young male p16-3MR mice displayed a clear induction of bioluminescent signals compared to baseline (Fig. 1A). Similarly, young female mice treated with doxorubicin (5 mg/kg for three consecutive days) exhibited a robust increase in luminescence two weeks later, which was absent in vehicle-treated controls (Fig. 1B). These findings demonstrate that the system can reliably capture the transient induction of senescence in response to injury and genotoxic stress.

**Figure 1.**
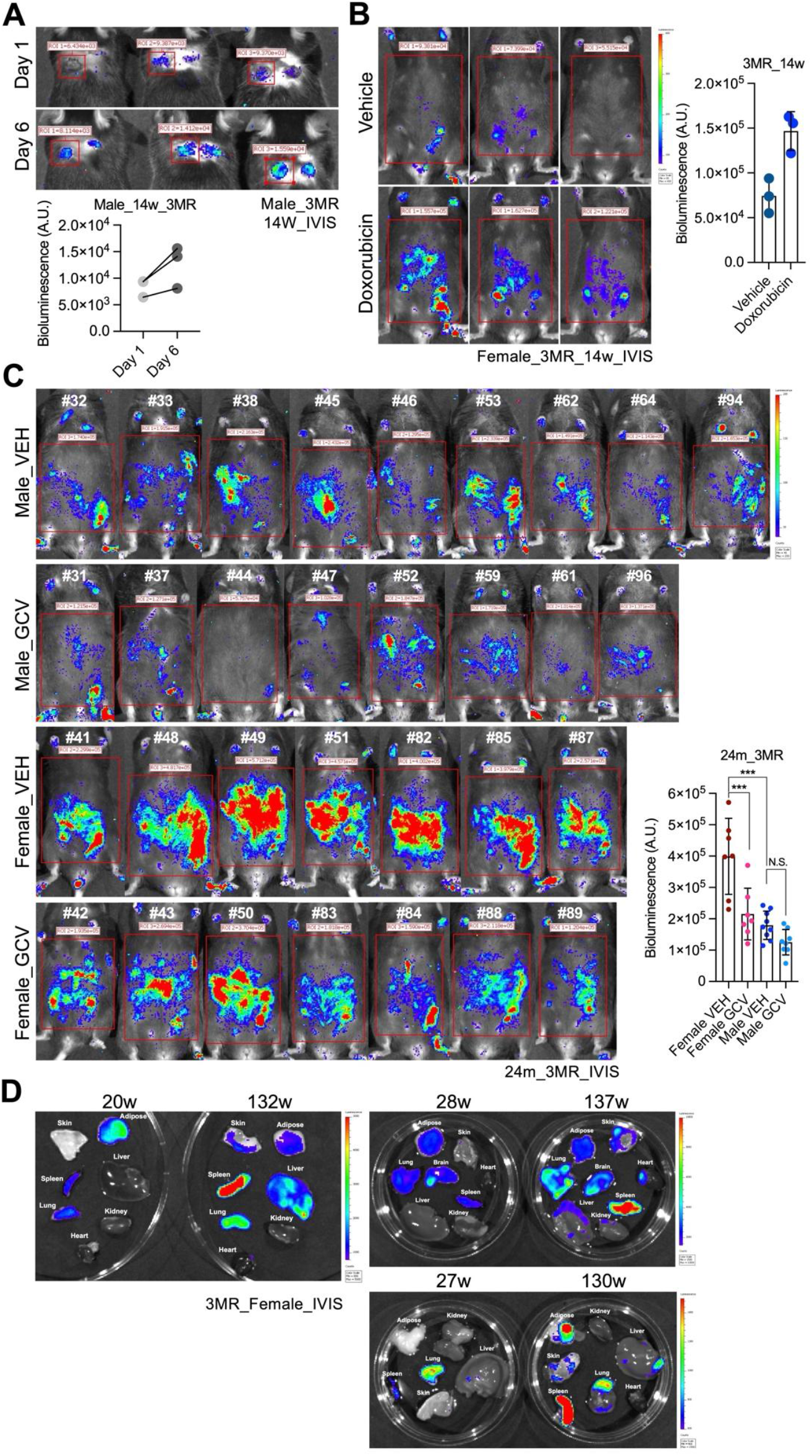
Validation of bioluminescence imaging for p16-3MR mouse model using IVIS Spectrum system. **(A)**Young male p16-3MR mice (∼14 weeks) were wounded with a 6-mm biopsy punch at day 0, injected intraperitoneally with coelenterazine-h, and imaged with the IVIS Spectrum system at days 1 and 6 post-wounding (n=3 mice/group; super binning; 5-min exposure). **(B)**Young female p16-3MR mice (∼12 weeks) treated with vehicle or doxorubicin (5 mg/kg for 3 consecutive days) were imaged 14 days later using IVIS Spectrum after coelenterazine-h injection (n=3 mice/group; super binning; 5-min exposure). **(C)**Male and female p16-3MR mice were treated monthly from 16 months of age with vehicle (pH 11 water) or ganciclovir (25 mg/kg in pH 11 water, 5 injections/month). At ∼24 months, mice were injected with coelenterazine-h and imaged using IVIS Spectrum. Groups were imaged together in random order (n=7–9 mice/group; super binning; 5-min exposure). **(D)***Ex vivo* imaging of organs from three independent pairs of young and old female p16-3MR mice. Animals were terminated in parallel, organs excised, incubated in 10× diluted coelenterazine-h (PBS) for 45–60 min, and imaged with IVIS Spectrum (5-min exposure; super binning).

We next assessed bioluminescent activity during aging and its modulation by clearance. Male and female p16-3MR mice were treated monthly with either vehicle or ganciclovir (GCV) starting at 16 months of age, and imaged at 24 months. Old females showed a higher baseline burden of bioluminescent signal compared to males, and this difference was significantly reduced following GCV treatment (Fig. 1C). Signal reduction was less pronounced in males, likely reflecting their lower baseline levels^26^. Importantly, to avoid bias, all animals were randomized and imaged together across treatment groups. These results support both the sex-specific dynamics of senescent cell accumulation and the ability of the HSV-tk/GCV system to achieve partial clearance *in vivo*.

To rule out the possibility that signals arose from non-specific sources such as abdominal fluid, fur, or skin pigmentation, we excised organs from matched pairs of young and old mice at different time points, incubated them in diluted coelenterazine-h, and performed *ex vivo* imaging. Old animals consistently displayed stronger bioluminescent signals than young animals across multiple tissues, including skin, spleen, liver, and lung (Fig. 1D). These findings confirm that the signals originate from tissues themselves and are associated with age. Comparable increases in signal have also been reported in other senescence-inducing settings using p16-3MR mice, including sublethal irradiation^27^ and cigarette smoke exposure^28^. Moreover, our results are consistent with findings from an independent p16-luciferase mouse model driven by a human promoter^8^, reinforcing the generalizability of luminescence-based detection of senescence.

Because Renilla luciferase catalyzes coelenterazine-h at an optimal emission wavelength near 480 nm^29–31^, we used the LAGO-X imaging system, which allows spectral filtering, as an additional and conservative validation of signal specificity. In shaved black-furred mice, open-filter imaging revealed substantial background signal, consistent with previous reports^11^. However, when a 490 nm filter was applied, this background was eliminated: young mice showed no detectable signal, whereas old mice retained a residual signal consistent with authentic Renilla luciferase activity (Fig. 2A). No signal was observed at irrelevant wavelengths (e.g., 870 nm), further confirming specificity.

**Figure 2.**
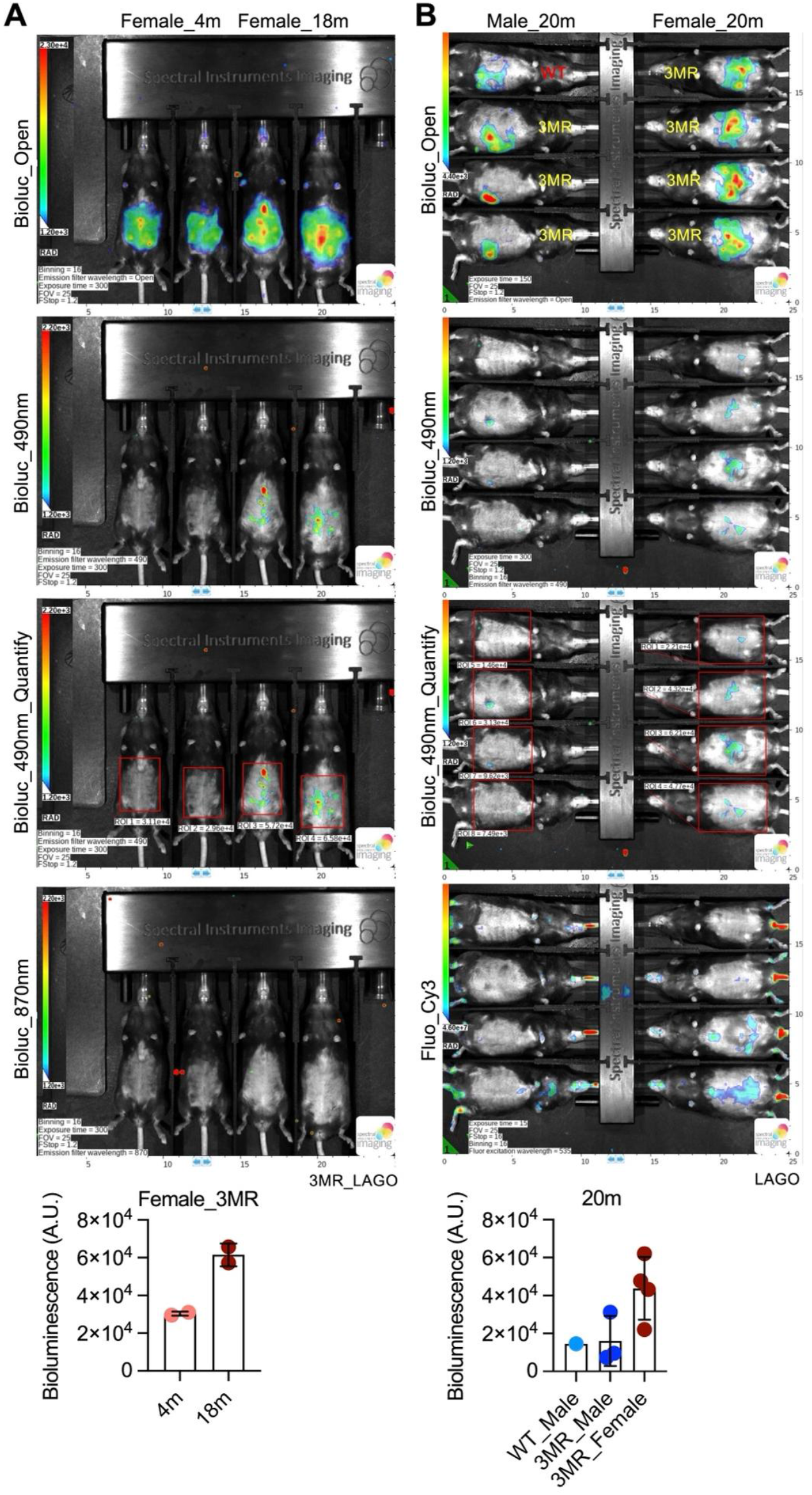
Validation of bioluminescence imaging for p16-3MR mouse model using LAGO-X system. **(A)**Young (∼4 months) and old (∼18 months) female p16-3MR mice were shaved and imaged with the LAGO-X system using open, 490-nm, or 870-nm emission filters. Signal from the 490-nm filter was quantified (n=2 mice/group; super binning; 5-min exposure). **(B)**Old male and female p16-3MR mice (∼20 months) were shaved and imaged with LAGO-X using open or 490-nm filters, or by fluorescence imaging with a Cy3 filter. One WT mouse was included as control (n=3–4 mice/group; super binning; 5-min exposure).

Finally, because we observed sex-related differences in signal intensity *in vivo* (Fig. 1C), we tested whether these persisted under filtered conditions and included WT controls. Non-specific background from coelenterazine-h was present in WT mice but was fully removed by 490 nm filtering (Fig. 2B). In p16-3MR mice, the sex disparity remained evident, with old females showing stronger signals than males (Fig. 2B), in accordance with a recent study from our group^26^. We also explored the utility of the RFP cassette for live imaging beyond flow cytometry. Consistent with other reports, RFP signals were weak and context-dependent, but again appeared stronger in females (Fig. 2B). Although the specificity of these signals requires further validation, these results underscore both the limitations and the potential complementary utility of the fluorescent component.

Together, these validation experiments confirm that the p16-3MR system is functional and reproducible across multiple senescence-inducing contexts, that these signals increase with age, and that they are partially reduced by GCV treatment. They also illustrate the known limitations of Renilla luciferase, such as background luminescence and reduced sensitivity in pigmented animals, which can be managed with proper filtering and controls.

## Discussion

The p16-3MR mouse model was developed to address a central challenge in senescence biology: the need to both identify and functionally interrogate p16-expressing cells *in vivo*. Over the past decade, its broad adoption across independent laboratories and biological contexts has generated a substantial body of evidence supporting its utility. The results presented here, together with extensive published work, reinforce several key conclusions regarding the strengths, limitations, and appropriate use of the p16-3MR system.

First, our findings confirm that the p16-3MR reporter is reproducibly induced in conditions associated with increased senescent cell burden, including tissue injury, genotoxic stress, and aging. These observations are consistent with numerous independent studies reporting p16-driven reporter activation in diverse tissues and disease settings. Importantly, these effects are not restricted to a single laboratory or experimental setup, underscoring that reporter induction reflects conserved biological processes rather than technical artifacts. The concordance between whole-body imaging, *ex vivo* organ analysis, and results obtained in other senescence models, including a human p16-driven luciferase reporter, further supports the conclusion that p16-linked luminescence tracks states when senescent cells are present in sufficient abundance.

Second, our data highlight the context dependence inherent to Renilla luciferase–based imaging. Compared with Firefly luciferase, Renilla luciferase provides lower photon output and is more sensitive to substrate biodistribution, pigmentation, and tissue depth. These properties limit detection sensitivity, particularly when senescent cells are sparse or localized deep within tissues. However, such constraints are intrinsic to the reporter chemistry and not specific to the p16-3MR construct. When senescent cell burden is high, as in aged tissues, damaged organs, or regenerating wounds, bioluminescent signals are robust and reproducible, even in pigmented animals. Spectral filtering, as employed here, offers an additional approach to distinguish authentic Renilla emission from nonspecific background, but is not a prerequisite for most applications reported in the literature. Together, these observations emphasize that interpretation of luminescence data should be guided by biological modulation and signal-to-background relationships rather than absolute photon counts.

Third, our results reaffirm the functionality of the HSV-tk/ganciclovir clearance module. Partial elimination of p16-expressing cells following ganciclovir treatment is consistently observed *in vivo* and aligns with prior reports across multiple tissues and disease models. Clearance efficiencies in the range of 50–75%, as observed here, are typical of HSV-tk–based ablation systems and reflect known constraints such as drug penetration, heterogeneous transgene expression, and bystander effects. Crucially, in the vast majority of published studies, senescent cell clearance in p16-3MR mice has been validated using orthogonal markers (primarily SA-β-gal activity, DNA damage markers, cell-cycle inhibitors, and SASP components) independent of the transgene itself.

Fourth, the differential use of the three components of the p16-3MR construct across studies is informative. As in other multi-module reporter systems, investigators have relied on HSV-tk for functional ablation, luciferase for longitudinal *in vivo* tracking, and mRFP primarily for *ex vivo* or flow-based applications. The more limited use of the fluorescent component reflects well-recognized constraints of mRFP brightness and tissue penetration, not a failure of the transgene. Indeed, similar usage patterns are observed in other senescence reporter models, underscoring that selective deployment of individual modules is a common and rational strategy.

More broadly, our findings illustrate a general principle in the use of complex genetic models: technical performance must be interpreted in light of biological context, experimental design, and validation strategy. Variables such as animal age, sex, genetic background, housing conditions, and imaging platform can all influence signal intensity, yet reproducibility across laboratories is best assessed by consistency of biological trends and validation experiments. The extensive independent literature using p16-3MR mice demonstrates that, when applied with appropriate controls and expectations, the model yields reproducible and biologically meaningful insights into senescent cell function.

Looking ahead, the field of senescence research will undoubtedly benefit from continued development of refined tools, including brighter reporters, alternative regulatory elements, and strategies to target specific senescent subpopulations. Such advances should be viewed as complementary to, rather than replacements for, existing models. The p16-3MR system has played a foundational role in establishing causal links between senescent cells and tissue dysfunction *in vivo*, and continues to serve as a benchmark against which newer approaches can be evaluated.

In summary, the results presented here, together with a decade of independent studies, demonstrate that the p16-3MR mouse model is functional, reproducible, and informative when used within its known technical boundaries. Rather than undermining its value, careful consideration of these boundaries strengthens experimental design and interpretation, ensuring that p16-3MR remains a robust tool for advancing our understanding of cellular senescence in physiology and disease.

## Materials & Methods

All the animal experiments were conducted in the Central Animal Facility (CDP) of the University Medical Center Groningen (UMCG) under conventional housing conditions. All experiments were approved by the Central Authority for Scientific Procedures on Animals in the Netherlands with the license numbers: AVD105002015339, AVD1050020184807 and AVD10500202115445. The p16-3MR transgenic mice and wild-type littermates were used to investigate the outcome of bioluminescence imaging in different experimental settings. For *in vivo* imaging, all the mice were injected intraperitoneally with 100 µL of Xenolight RediJect Coelenterazine h (PerkinElmer, 760506). Twenty minutes post-injection, mice were anesthetized with 2% isoflurane, and Renilla luciferase bioluminescence was captured using the IVIS Spectrum *in vivo* Imaging System (PerkinElmer) or LAGO-X Spectral Instruments Imaging with super binning (5 min exposure) at CDP-UMCG. For wound healing experiment, young male p16-3MR mice (∼14 weeks old) were wounded with a 6-mm biopsy punch at day 0, and imaged with the IVIS Spectrum System at days 1 and 6 post-wounding (n=3 mice/group). For chemotherapy experiment, young female p16-3MR mice (∼12 weeks old) were treated with vehicle or doxorubicin (5 mg/kg for 3 consecutive days) and then imaged 14 days later using the IVIS Spectrum System. For the natural aging experiment, both male and female p16-3MR mice were treated monthly from 16 months old with vehicle (pH 11 water) or ganciclovir (25 mg/kg in pH 11 water, 5 injections/month). At around 24 months old, mice were imaged using IVIS Spectrum System. Mice from different groups were imaged together in random order (n=7–9 mice/group). For the *ex vivo* imaging, the organs from three independent pairs of young and old female p16-3MR mice were imaged at different time-points. Animals were terminated in parallel, organs were excised and incubated in 10 times diluted coelenterazine-h (PBS) for 45–60 min and imaged with the IVIS Spectrum System. For the fur shaving and bioluminescence filtering experiments in young and old mice, LAGO-X was used to apply the open, 490-nm, or 870-nm emission filters and image the mice.

## Data availability

No new dataset was generated in this study. Reasonable requests for materials can be made directly to the corresponding authors.

## Acknowledgements

The project was funded by grants to the laboratory of M.D. from the Nederlandse Organisatie voor Wetenschappelijk Onderzoek, the Dutch Cancer Foundation (KWF) and the Hevolution Foundation.

## Author contributions

B.W. and M.D. conceptualized the study. B.W., J.N., and M.D. devised the study methodology.

B.W. and J.N. conducted the investigation. B.W., and J.N. visualized data. M.D. acquired the funding and managed the project. B.W. and M.D. supervised the study. B.W. and M.D. wrote the original manuscript draft. B.W., P.P.R,, J.N., and M.D. reviewed and edited the manuscript.

### Box 1.

**Practical considerations for using p16-3MR mice**

1. Controls: Use littermate WT controls where feasible; avoid commercially sourced WT animals when assessing low-level signals.
2. Imaging: Prioritize biological modulation over absolute photon counts; compare conditions within the same imaging session.
3. Pigmentation: Shaving may improve sensitivity but introduces biological confounds; interpret cutaneous signals cautiously.
4. Validation: Always corroborate findings with senescence markers independent of the transgene.
5. GCV dosing: Use established regimens (25–50 mg/kg, short duration); avoid extrapolating effects from high-dose antiviral studies.
6. Interpretation: Expect partial clearance and context-dependent outcomes.

